# Tartrate-resistant acid phosphatase (TRAP/ACP5) sex-specifically regulates bone maintenance in old mice, but the anabolic effects of mechanical loading is regulated in a sex-independent way

**DOI:** 10.1101/2025.09.03.673915

**Authors:** Bhavik Rathod, Jasmine Samvelyan, Nicole Gustafsson, Aneta Liszka, Narelle McGregor, Jianyao Wu, Claes Ohlsson, Anna Fahlgren, Natalie Sims, Jonas Fuxe, Göran Andersson, Jessica J Alm, Sara H Windahl

**Author notes:** Shared last authorship (Jessica J Alm, Sara H Windahl). **Corresponding author:** Sara Windahl, Associate Professor, Ph.D. Karolinska Institutet. ANA Futura, Department of Laboratory Medicine, Division of Pathology Alfred Nobels Allé 8, 141 52 Huddinge, Sweden.

## Abstract

Tartrate-resistant acid phosphatase (TRAP/ACP5), primarily known as an osteoclast marker, has emerged as a critical regulator of skeletal integrity, regulating sex-specific bone growth, and bone’s response to mechanical load in young adult male mice. In this study, we investigated the sex-specific roles of TRAP in bone structure and response to mechanical stimuli in old (19-month-old) wild-type (WT) and TRAP-deficient (TRAP^-/-^) mice using micro-computed tomography, serum bone turnover markers, *in vivo* axial mechanical loading, and *in vitro* mechanotransduction assays. Our findings revealed that TRAP^-/-^ mice of both sexes maintained shorter tibiae than WT mice independent of sex. Notably, male, but not female, TRAP^-/-^ mice have increased trabecular bone volume fraction and cortical bone area compared to WT, indicative of disrupted bone remodelling processes in male mice. Interestingly, TRAP-deficiency substantially impaired the anabolic bone response to mechanical loading, affecting both trabecular and cortical compartments in both sexes, indicating that when challenged, TRAP is important for bone formation also in female mice. Mechanical stimulation *in vitro* of hematopoietic progenitor cells from WT and TRAP^-/-^ mice revealed that the increased ATP-release in response to mechanical stimulation was only disrupted in male mice, while mechanically induced increase in osteoclast formation was inhibited in TRAP^-/-^ mice of both sexes. These results highlight the importance of TRAP in maintaining trabecular architecture and cortical bone in male mice and underscore its critical function in mediating adaptive responses to mechanical loading of both sexes, during aging. Future investigations should focus on elucidation of TRAP-dependent pathways as potential therapeutic targets to counteract age-related deficits in bone adaptation and remodelling.

## Introduction

Bone strength and integrity depend on coordinated bone remodelling, regulated by mechanical loading and endocrine signalling, resulting in sex- and site-specific differences in skeletal phenotype. Bones adapt structurally via mechanotransduction, a mechanism primarily mediated by osteocytes sensing mechanical forces and signalling to osteoblasts and osteoclasts^1,2^. Tartrate-resistant acid phosphatase (TRAP), a recognized osteoclast marker, also performs crucial enzymatic functions, including dephosphorylation of bone matrix proteins such as osteopontin (OPN) and bone sialoprotein (BSP), facilitating osteoclast migration and bone matrix remodelling ^3,4^.

Our previous research also has demonstrated TRAP’s regulatory impact on sex- and site-specific skeletal responses^5^. We showed that young TRAP-deficient (TRAP^-/-^) mice have impaired bone remodelling characterized by delayed cartilage mineralization, widened and disorganized growth plates, shortened bones, mild osteopetrosis, and reduced response to mechanical loading ^4–7^. These skeletal changes display clear site-specificity, with pronounced abnormalities evident in weight-bearing bones such as the tibia, suggesting that mechanical loading significantly influences TRAP-dependent skeletal adaptation. Additionally, these effects differ notably by sex, with male TRAP^-/-^ mice exhibiting greater alterations in bone geometry and trabecular microarchitecture compared to females, while female TRAP^-/-^ mice display a more severe growth plate phenotype than males ^5,6^. Such observations emphasize the role of TRAP in regulating both site- and sex-specific bone development and remodelling, raising further questions about the underlying cellular and molecular mechanisms involved.

Mechanical loading regulates bone remodelling primarily through osteocytes, which coordinate the release of signalling molecules essential for bone adaptation. Mechanical stimulation induces osteocytes and osteoblasts to release extracellular ATP, which acts as a critical mediator in mechanotransduction. ATP activates purinergic (P2) receptors, initiating signalling cascades that promote osteogenic differentiation and bone remodelling—an effect clearly demonstrated by in vitro loading studies ^8^. Notably, recent evidence shows that TRAP deficiency impairs this adaptive bone formation response to mechanical loading in male mice, underscoring TRAP’s importance in mechanotransducive processes^7^. Therefore, the absence of TRAP could disrupt key osteocyte-driven signalling pathways involving both sclerostin and extracellular ATP, ultimately reducing bone’s capacity to effectively respond to mechanical demands^7,8^.

Aging affects bone remodelling by reducing osteocyte viability, impairing osteoblast function, enhancing osteoclast-driven bone resorption, and elevating chronic inflammation (inflammaging) ^9–11^. TRAP, through its modulation of inflammatory cytokines such as osteopontin, TNF-α, and IL-1β, could influence this inflammatory response^3,12,13^. Lifelong absence of TRAP might therefore amplify age-associated skeletal deterioration by intensifying inflammatory signals and altering immune cell functions. Human TRAP deficiency (spondyloenchondrodysplasia) manifests with combined skeletal and immune dysregulation, highlighting TRAP’s broader involvement in osteoimmunology^14–16^.

In the present study, we performed an in-depth sex- and site-specific characterization of bone in old (19-month-old) TRAP^-/-^ mice, in order to assess whether the observed alterations in bone structure and adaptive response to mechanical loading in young adult mice is maintained in old mice.

## 2. Materials and Methods

### 2.1 Animals

Mice heterozygous for TRAP were bred to generate male and female wild-type (WT) and TRAP^-/-^ offspring of both genotypes^5^. The mice were housed in a standard animal facility (KMB, Karolinska Institutet) in groups of up to five animals (both genotypes) per cage in IVC-500 cages under controlled conditions (22°C, 12-hour light/dark photoperiod). They were provided water and a pellet diet (CRM(P), Special Diet Services, Scanbur) for the initial 2 months, followed by CRM(P), Special Diet Services, SAFE, ad libitum. The animals were allowed to age to 18 months ± 2 weeks months before further analyses. All animal experiments were approved by the local Ethical Committees for Animal Research (ethical numbers 8387-2018 with amendments, 11055-23 and 19634-2023) and adhered to the ARRIVE guidelines. Animals were euthanized by blood extraction from auxiliary blood vessels under ketamine (Abcur) and dexdomitor (Orion Pharma AB) anaesthesia, followed by cervical dislocation. Genotyping was performed as previously described^5^. For NGS analysis 2-3 months old C57BL/6 male mice were used.

### 2.2 *Ex-Vivo* Strain Measurements

For controlled and comparable bone-loading set-ups, *ex-vivo* strain gauging analyses were performed on a subset of old WT and TRAP-/- mice to identify the correct applicable loading values for genotypes and sexes. The methodology for *ex-vivo* strain measurements of the mouse tibia has been detailed previously ^17^. In brief, the magnitude of axial mechanical strain applied to the tibia during loading was determined *ex vivo* in euthanized subgroups (N = 6–7 per experimental group) using a 3100 ElectroForce® Test Instrument (TA Instruments, MN, USA) to ensure consistent peak strain levels across groups. Peak compressive loads ranging from 6 to 18 N were applied to determine corresponding strain levels. The same ramping trapezoidal waveform was used for these loads, and the identical holding cups and loading apparatus were later employed for *in vivo* loading. Linear regression analysis was performed to evaluate the data.

### 2.3 Mechanical Loading

In this study, an established *in vivo* tibial loading protocol was utilized, as previously described^17^. Briefly, mice were positioned so that both the knee and ankle joints, maintained in flexion, were placed into concave cups. The upper cup (encompassing the knee) was connected to the loading device’s actuator arm, while the lower cup was attached to a dynamic load cell. This configuration ensured that the tibia remained under a continuous static preload of 0.5 N.

The protocol targeted a peak mechanical strain of 1800 με according to previously established loading-responses. Peak mechanical strain at 1800 microstrain (με) was achieved at 10.4 N in WT male mice, at 11.4 N for TRAP-/- male mice, and 9.4 N for both WT and TRAP-/- female mice (Supplementary 1 A). Subsequently, for the in vivo loading protocol, 40 cycles of dynamic loading were applied, each followed by a 10-second rest period at 0.5N. The loading session were performed 3 days a week for 2 weeks. All mice were euthanized three days after completion of the final loading session.

### 2.4 Serum Enzyme-Linked Immunosorbent Assay (ELISA)

Blood was collected from auxiliary blood vessels and transferred into Multivette® serum gel tubes. Serum was extracted following the manufacturer’s protocol. C-terminal telopeptide of type I collagen (CTX) (AC-06F1, IDS) and N-terminal propeptide of type I procollagen (PINP Rat/Mouse EIA AC-33F1, IDS) levels were measured following the manufacturer’s protocol.

### 2.5 Micro-Computed Tomography (μCT) of trabecular and cortical bone

Tibiae were carefully dissected to remove surrounding muscle tissue, fixed in 4% formalin for 48 hours, and then transferred to 70% ethanol for preservation. μCT analysis was performed using a Skyscan 1276 scanner (Skyscan N.V., Aartselaar, Belgium) with the following parameters: X-ray tube voltage of 40 kV, current of 200 μA, and a 0.25-mm aluminium filter. Scanning was performed with a 180° angular rotation and a 0.4° angular increment. The exposure time per step was 520 milliseconds, and the resulting voxel size was isotropic at 9 μm.

Reconstruction of the datasets was performed using NRecon software (version 2.2.0.6), followed by segmentation into binary images using adaptive local thresholding. Trabecular bone within the proximal tibia was selected for analysis. The region of interest (ROI) excluded cortical bone and began 3% of the total bone length distal to the growth plate, extending longitudinally for 0.271 mm (equivalent to 30 slices). Trabecular thickness and separation were quantified using the sphere-fitting local thickness method. Cortical bone analysis was done at 15% and 37% site for males and females respectively, where the ROI was selected extending longitudinally for 0.271 mm (equivalent to 30 slices).

### 2.6 Generation of mouse hematopoietic progenitor cells

Upon sacrifice, front legs were collected and transferred into sterile PBS solution and kept on ice for transport. Isolated bone marrow cells were cultured in incubation medium containing Minimum Essential Medium α (Thermo Fisher Scientific, Stockholm, Sweden) supplemented with 5% fetal bovine serum (Cytiva Europe GmbH Tyskland, Uppsala, Sweden), 1% antibiotic– antimycotic solution (Thermo Fisher Scientific) and 100 ng/ml recombinant mouse macrophage colony-stimulating factor protein (R&D Systems, Minneapolis, MN). After 48 h, these adherent cells were washed with phosphate-buffered saline (PBS, pH = 7.2, Life Technologies Europe BV), harvested with ice-cold 0.02% EDTA solution (Merck), and used for further experiments

### 2.7 *In vitro* model for mechanical stimulation

Mouse hematopoietic progenitor cells were seeded onto 5 μg/cm^2^ bovine Fibronectin-coated glass slides (Merck) at a density of 100.000 cells/cm^2^ 24 h prior to mechanical loading. Cells were subjected to pulsating fluid flow (PFF) using a parallel-plate flow chamber. A detailed characterization of these different loading profiles has been described previously (High shear stress amplitude in combination with prolonged stimulus duration determine induction of osteoclast formation by hematopoietic progenitor cells ^18^. Briefly, low intensity loading (Low) (0.7 ± 0.3 Pa, 5 Hz) was applied as a sinusoidal wave to reassemble normal loading in the bone ^19^. While supraphysiological loading (3.0 ± 0.2 Pa, 1Hz) was applied as a square wave, to reassemble high pressurized fluid flow (High). The non-loaded control (NL) was used to represent a lack of mechanical stimulation on the cells. Eight millilitres of serum-free fluid flow medium that consisted of MEMα with 1% PSF was used. Conditioned medium after 2 minutes of PFF was collected and either snap frozen in liquid nitrogen and stored at −80°C (ATP and LDH measurement) or sterile filtered and frozen at −20°C (Osteoclastogenesis assay).

### 2.8 RANKL-induced osteoclastogenesis assay

Whole bone marrow of two wild-type males C57BL/6 mice (9–12 weeks, Janvier Labs) was pooled and seeded in 96-well plates at a density of 3.125 × 10^5^ cells/cm^2^. For each condition, triplicates were performed. The culture medium was a mixture of 50% freshly prepared medium and 50% conditioned medium. Each well contained αMEM (Life Technologies) with 5% FBS (Biowest) and 1% PSF (Life Technologies EuropeBV) supplemented with 30 ng/mL recombinant murine M-CSF (R&D systems, Inc.) and 20 ng/mL recombinant murine RANKL (R&D systems, Inc.). A positive control was performed with M-CSF and RANKL suspended in the corresponding volumes with fresh medium, while in the negative control, RANKL was omitted. After 3 days, the medium was changed and after 6 days the cells were fixed with 4% formaldehyde and stained for tartrate-resist-ant acid phosphatase (TRAP) according to the manufacturer’s instructions (Sigma-Aldrich Sweden AB). The TRAP-positive multinucleated cells (≥3 nuclei/cell) were counted manually using an Axio Vert.A1 fluorescence microscope (Carl Zeiss AB) with an N-Achroplan 20×/0.45 M27objective (Carl Zeiss AB)

### 2.9 ATP measurements

Extracellular ATP was measured using the ATP determination kit (Fisher Scientific). Assay was performed according to manufactures instructions with snap frozen conditioned medium after 2 minutes of mechanical loading.

### 2.10 Statistical analyses

Data are presented as mean ± standard deviations, alongside individual data points. All data were tested for normal distribution and equal variances using Kolmigorov-Smirnov test and Levene’s test, respectively. Data fulfilling these assumptions were analyzed using parametric statistics. For analyzing the effect of TRAP on phenotypic parameters, a two-way ANOVA was conducted to determine to what extent genotype and sex have effects on the parameters, and possible interactions. For analyzing the effect of mechanical loading, a repeated-measures two-way ANOVA was conducted. The primary focus was on intrasexual effects of TRAP, and secondary focus was on sex-related differences within the same genotype. In cases with a significant interaction, the main effects were interpreted jointly. In cases without significant interaction, the interaction was removed from the model and analyses were re-run without the interaction term. Tukey’s post hoc test, T-test, or paired T-test were applied for pairwise comparisons, depending on the data and specific question. Statistical significance was defined as p < 0.05.

## 3. Results

### 3.1 Distinct effects of TRAP on body weight and internal-organ size in old female but not male mice

To investigate if TRAP deficiency alters somatic growth or internal-organ size in old male and female mice, we assessed body and organ weights of 18-month-old TRAP-knockout (TRAP^-/-^) mice and their wild-type (WT) littermates. The mice displayed pronounced sexual dimorphism in terms of effects of deleting TRAP, with significant alterations in female TRAP^-/-^ mice, while body weight, internal-organ sizes and blood glucose levels were unaltered in male TRAP^-/-^ mice compared to their WT littermates, respectively (Table 1). The TRAP^-/-^ females had a significant −20% reduction in body weight compared to WT littermates, whereas TRAP^-/-^ males showed a mean reduction of 11% in body weight compared to WT males, which was not statistically significant (Table 1). Due to the significant differences in body weight, the internal-organ sizes were normalized to body weight for further comparisons between genotypes and sexes. Female TRAP^-/-^ mice had a −28% reduction in gonadal fat, a +98% increase in uterus, and a +30% increase in lung weights, respectively, compared to WT females. No differences were detected in spleen, liver, thymus or in blood glucose levels in female TRAP^-/-^ compared to WT littermates.

**Table 1.** Body mass and internal-organ sizes in WT and TRAPK -/- male and female mice 20-weeks of age.

|  |  |  |  |  |  |  |  |  |
| --- | --- | --- | --- | --- | --- | --- | --- | --- |
|  | Males |  |  | Females |  |  | WT | TRAP -/- |
| Body mass (BM) and Organ mass | WT | TRAP -/- | p-value | WT | TRAP -/- | p-value | p-value male vs female | p-value male vs female |
| Body mass (g) | 41.1 ± 8.6 | 36.7 ± 7.8 | 0.350 | 55.8 ± 4.5 | 44.7 ± 11.5 | **0.022** | **0.0004** | 0.194 |
| Gonadal fat (mg) / BM | 0.03 ± 0.02 | 0.03 ± 0.01 | 0.793 | 0.13 ± 0.03 | 0.09 ± 0.04 | **0.049** | **<0.0001** | **0.008** |
| Spleen (mg) / BM | 0.002 ± 0.001 | 0.003 ± 0.001 | 0.119 | 0.003± 0.001 | 0.004± 0.001 | 0.263 | 0.057 | 0.104 |
| Lung (mg) / BM | 0.005 ± 0.001 | 0.005 ± 0.001 | 0.587 | 0.003 ± 0.0001 | 0.004 ± 0.001 | **0.047** | **0.001** | **0.034** |
| Thymus (mg) / BM | 0.001 ± 0.001 | 0.001 ± 0.001 | 0.836 | 0.002± 0.001 | 0.001± 0.001 | 0.225 | **0.038** | 0.355 |
| Liver (g) / BM | 0.05± 0.007 | 0.05 ± 0.007 | 0.906 | 0.04± 0.01 | 0.04± 0.01 | 0.101 | **0.016** | **0.0008** |
| Seminal vesicles (mg) or Uterus (mg) / BM | 0.04 ± 0.03 | 0.03 ± 0.02 | 0.507 | 0.003 ± 0.002 | 0.006 ± 0.003 | **0.023** | NA | NA |

Data presented as mean +/- SD

When further investigating sex-differences in body mass and organ weights in old mice, clear differences were detected between males and females in all parameters. In WT mice, females displayed +36% higher body mass, +27% higher spleen weight, +58% higher thymus weight, - 37% lower lung weight, and −18% lower liver weight, compared to WT males. The most pronounced difference was in the gonadal fat, where WT females had 3.8-times more than WT males (+285%). Less pronounced sex-differences were detected in TRAP^-/-^ mice, with only differences being a 3-times higher gonadal fat (+200%), −22% lower lung weight and −24% lower liver weight in TRAP^-/-^ females compared to TRAP^-/-^ males.

### 3.2 Shorter tibial length in TRAP*-/-* mice independent of sex

To investigate whether TRAP exerts sexually dimorphic effects on bone length during aging, we compared tibial length in male and female WT and TRAP*-/-* mice. For tibial length, there was an effect of genotype, but not sex, with TRAP^-/-^ mice exhibiting significantly reduced tibial length compared to their WT counterparts (Fig. 1A). Post hoc analysis revealed similar reduction in tibial length, of −13% in TRAP^-/-^ males and −12% in TRAP-/- females compared to their WT littermates. To further investigate bone remodelling status in old mice, circulating bone turnover markers (CTX-1 and P1NP) were measured. These markers primarily reflect systemic bone turnover rather than mechanisms directly regulating bone length, and their levels are known to be lower in old animals compared to young ones. Both male and female TRAP^-/-^ mice showed significant reduction in serum levels of resorption marker CTX, with a −6.4% reduction in male TRAP^-/-^ mice, and a −5.6% reduction in female TRAP^-/-^ mice, compared to their WT littermates (Fig 1 B). In contrast, serum levels of formation marker P1NP was similar in TRAP^-/-^ and WT male and female mice (Fig 1C). No sex-differences were detected in tibial length, or serum markers CTX and P1NP.

**Figure 1.**
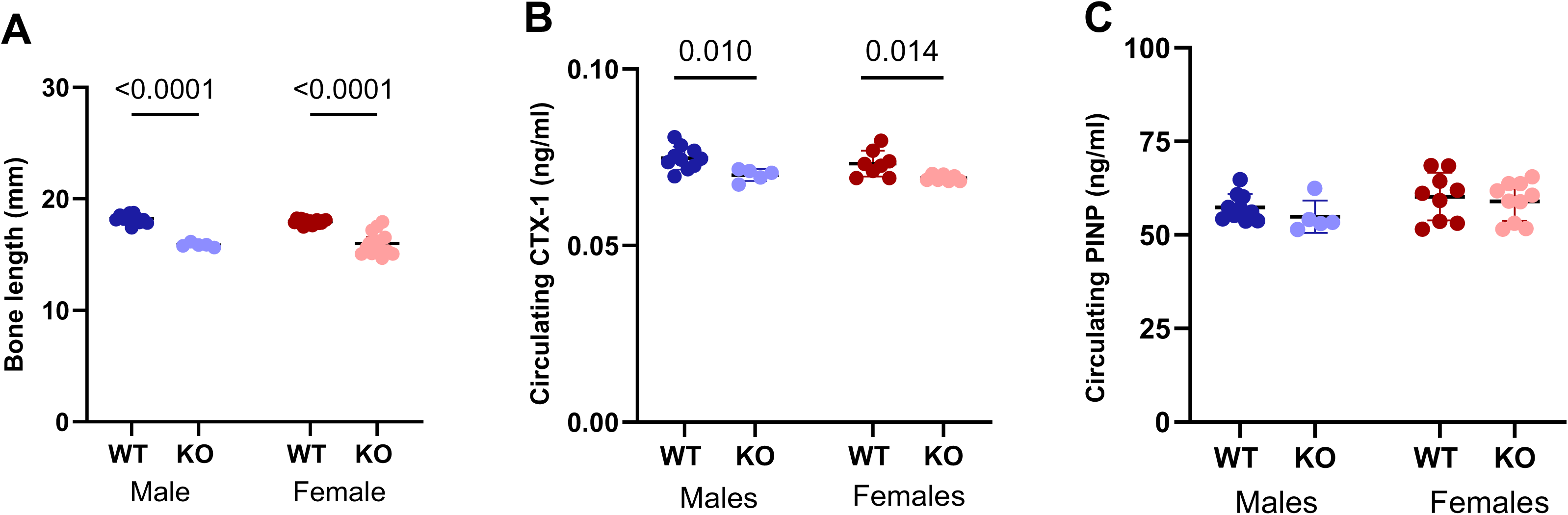
Shorter tibial length in TRAP^-/-^ mice. A) Tibial length, B) serum C-terminal telopeptide of type I collagen (CTX-1) levels, and C) serum procollagen type I N-terminal propeptide (P1NP) levels in WT and TRAP^-/-^ (KO), male and female mice. Data are presented as individual points with mean ± SD. Two-way ANOVA followed by Tukey’s multiple comparison test with *p*-values for WT vs-KO indicated in the graph. *N* = 5–10 per sex and genotype.

### 3.3 Sexually dimorphic effects of TRAP^-/-^ in old cortical bone

To assess the impact of TRAP deficiency and cortical bone morphology, total cross-sectional area (Tt.Ar), cortical area (Ct.Ar), marrow area (Ma.Ar), and cortical bone mineral density (Ct.BMD) of the tibia were measured in male and female WT and TRAP^-/-^ mice. These analyses showed different patterns of cortical bone effects of TRAP in males and females. Male TRAP^-/-^ mice exhibited a significant +26% increase in total cross-sectional area and a +30% increase in cortical area compared to WT males (Fig. 2A-B), while no differences were detected in marrow area (Fig. 2C) or cortical BMD (Fig. 2D). Female mice demonstrated a completely different pattern of effects of TRAP on cortical bone. Female TRAP^-/-^ mice displayed no differences in total cross-sectional area or cortical area compared to WT females (Fig. 2A-B), while the marrow area was significantly reduced by −32%, and the cortical BMD increased by +16% (Fig. 2C-D).

**Figure 2.**
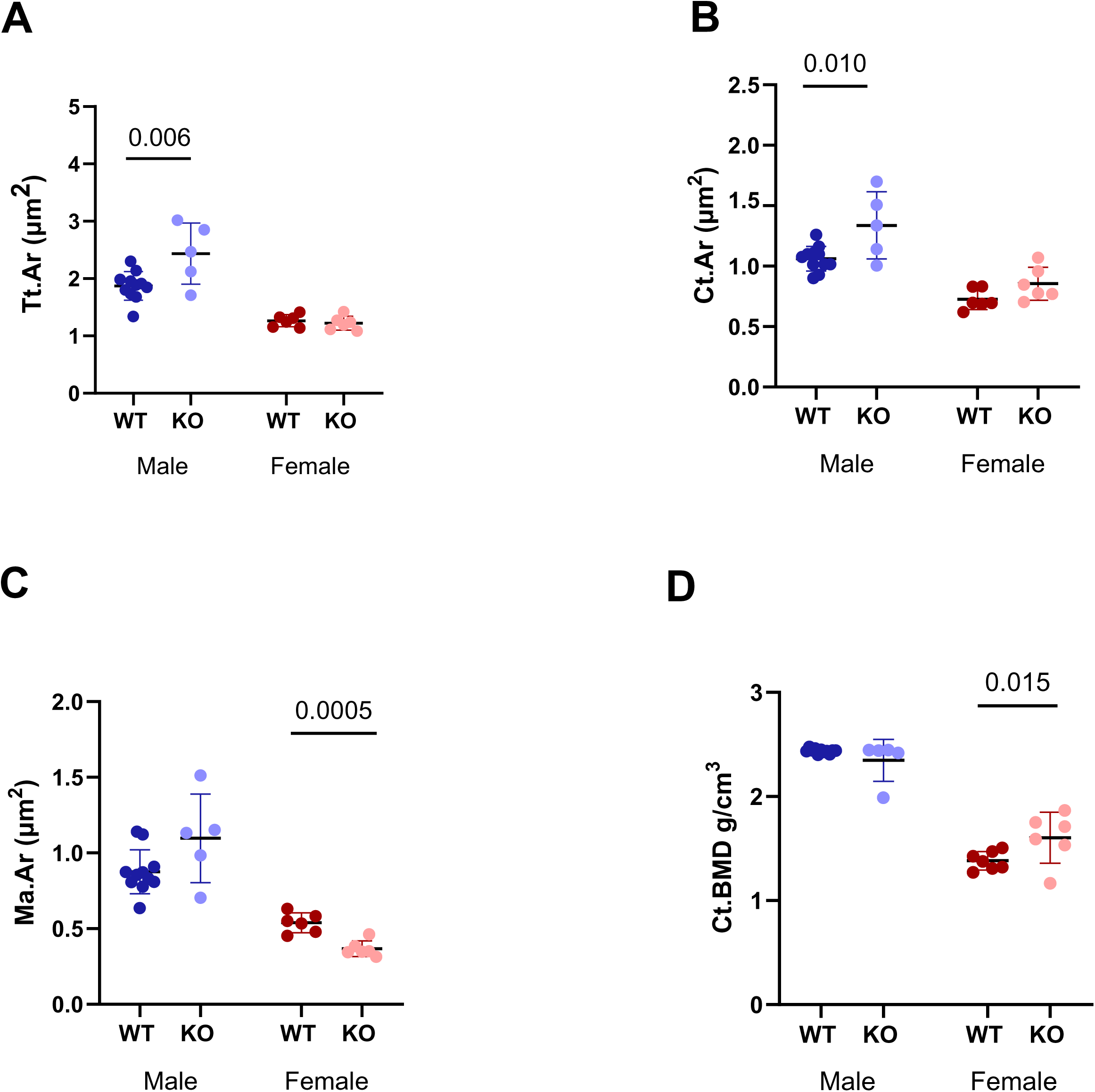
Sexually dimorphic effects of TRAP on cortical bone in old mice. Micro-computed tomography (μCT) analyses of tibiae from wild-type (WT) and TRAP^-/-^ (KO) male and female mice. A) Total cross-sectional area (Tt.Ar), B) cortical area (Ct.Ar) and C) marrow area (Ma.Ar), and D) cortical BMD. Data are presented as individual points with mean ± SD. Two-way ANOVA followed by Tukey’s multiple comparison test with *p*-values for WT vs-KO indicated in the graph. *N* = 5–10 per sex and genotype.

Comparisons between males and females further demonstrated the sexual dimorphisms in cortical bone measurements. In female WT mice, all the cortical bone parameters were significantly lower compared to WT males. Both cross-sectional area and the cortical area, were −32% lower (p=0.002 and p<0.0001, respectively)(Fig. 2A-B), while the marrow area was −39% lower (p=0.002) in female compared to male WT mice (Fig. 2C). The most pronounced difference was seen in cortical BMD, which was −43% lower in females compared to males (<0.0001)(Fig. 2D). The sex-differences were even more pronounced when comparing TRAP^-/-^ females with TRAP^-/-^ males. TRAP^-/-^ females had −50% lower total area (p<0.0001), −36% lower cortical area (p=0.004), −66% lower marrow area (p<0.0001), and −32% lower cortical BMD (p<0.0001). This probably reflecting, and amplifying, the large differences already present between WT male and female old cortical bone, in combination with the observed increases in TRAP^-/-^ males, and lack of response or decrease in TRAP^-/-^ females.

### 3.4 Pronounced effects of TRAP-/- on old trabecular bone in males, but not in females

To investigate the impact of TRAP deficiency on old trabecular bone maintenance, tibial trabecular bone was analysed by μCT in WT and TRAP^-/-^ mice. Analyses revealed clear sex-specific effects of TRAP on trabecular bone parameters, with significant alterations in TRAP^-/-^ males, while trabecular bone was normal in TRAP^-/-^ females, compared to their WT counterparts (Fig. 3). Compared to WT males, TRAP^-/-^ male mice had significantly increased trabecular bone, as demonstrated by more than two times higher (+121%) trabecular bone volume fraction (BV/TV)(Fig. 3A), +26% higher trabecular thickness (Tb.Th)(Fig. 3B), and two times higher (+102%) trabecular number (Tb.N)(Fig. 3C). No differences were detected in trabecular separation (Tb.Sp)(Fig. 3D).

**Figure 3.**
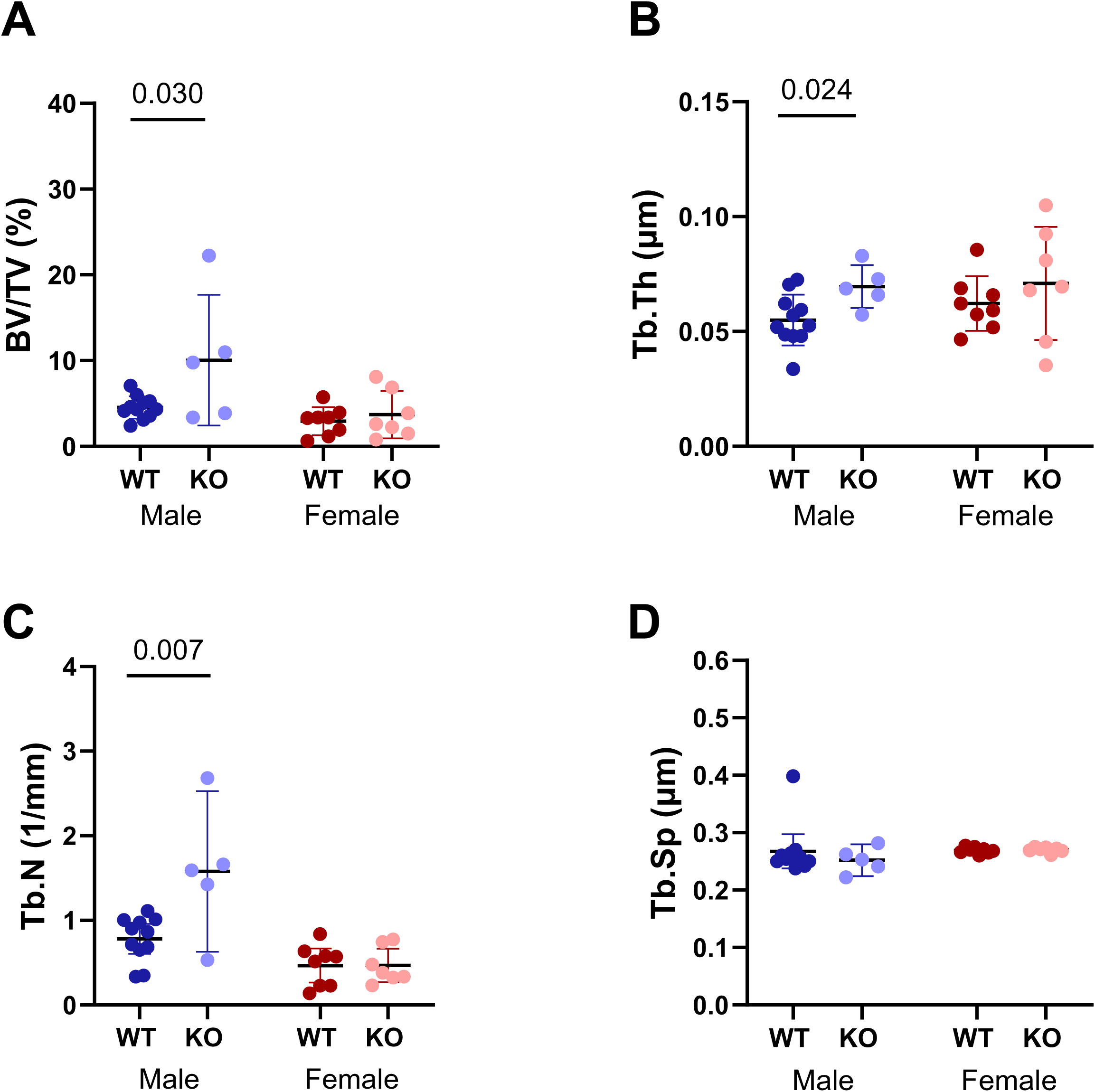
Pronounced effects of TRAP on old trabecular bone in male mice. Micro-computed tomography (μCT) analyses of tibiae from wild-type (WT) and TRAP^-/-^ (KO) male and female mice. (A) trabecular bone volume fraction (BV/TV), (B) trabecular thickness (Tb.Th), (C) trabecular number (Tb.N), and (E) trabecular separation (Tb.Sp). Data are presented as individual points with mean ± SD. Two-way ANOVA followed by Tukey’s multiple comparison test with *p*-values for WT vs-KO indicated in the graph. *N* = 5–10 per sex and genotype.

To investigate sex-differences in trabecular bone parameters in old mice, sub analyses were conducted and revealed significantly lower trabecular bone in female compared to male WT mice. The female mice had a −35% lower trabecular bone volume fraction (p=0.030), −40% lower trabecular number (p=0.016) and a +5.8% increased trabecular separation (p=0.002), while no difference was detected in trabecular thickness (Tb.Th) (Fig. 3A-D).

### 3.5 Reduced growth plate bony bridges in old female TRAP^-/-^ mice

We have previously shown reduced growth plate bony bridges in young adult male TRAP^-/-^ mice, without any detected effects in young adult female TRAP^-/-^ mice ^5^. We therefore investigated the number and areal density of growth plate bony bridges in old TRAP^-/-^ mice. As with other bone parameters, there were clear sex-specific differences in the effect of TRAP also on the growth plate bony bridges. For old female TRAP^-/-^ mice, significant reductions were seen in number and areal densities of bony bridges (Fig. 4). The total number of epiphyseal growth plate bony bridges was reduced by −37% in TRAP^-/-^ females compared with WT females (Fig. 4A). Further analyses revealed that this was due to the significant −36% reduction in number of medial and −43% reduction in lateral bridges, respectively (Fig. 4A-C). Similarly, the areal density of bony bridges was significantly reduced by −27% in the total epiphyseal area, reduced by −26% in the medial epiphyseal area, and reduced by −29% in the lateral epiphyseal area in old TRAP^-/-^ females compared to their WT littermates (Fig. 4D-F). In contrast, effects were much milder in males, with the only detected effects seen in the lateral tibiae, with a −33% reduction in number of bridges and a −26% reduction in areal bridge density in old TRPKO male mice compared to WT males (Fig 4C, F).

**Figure 4.**
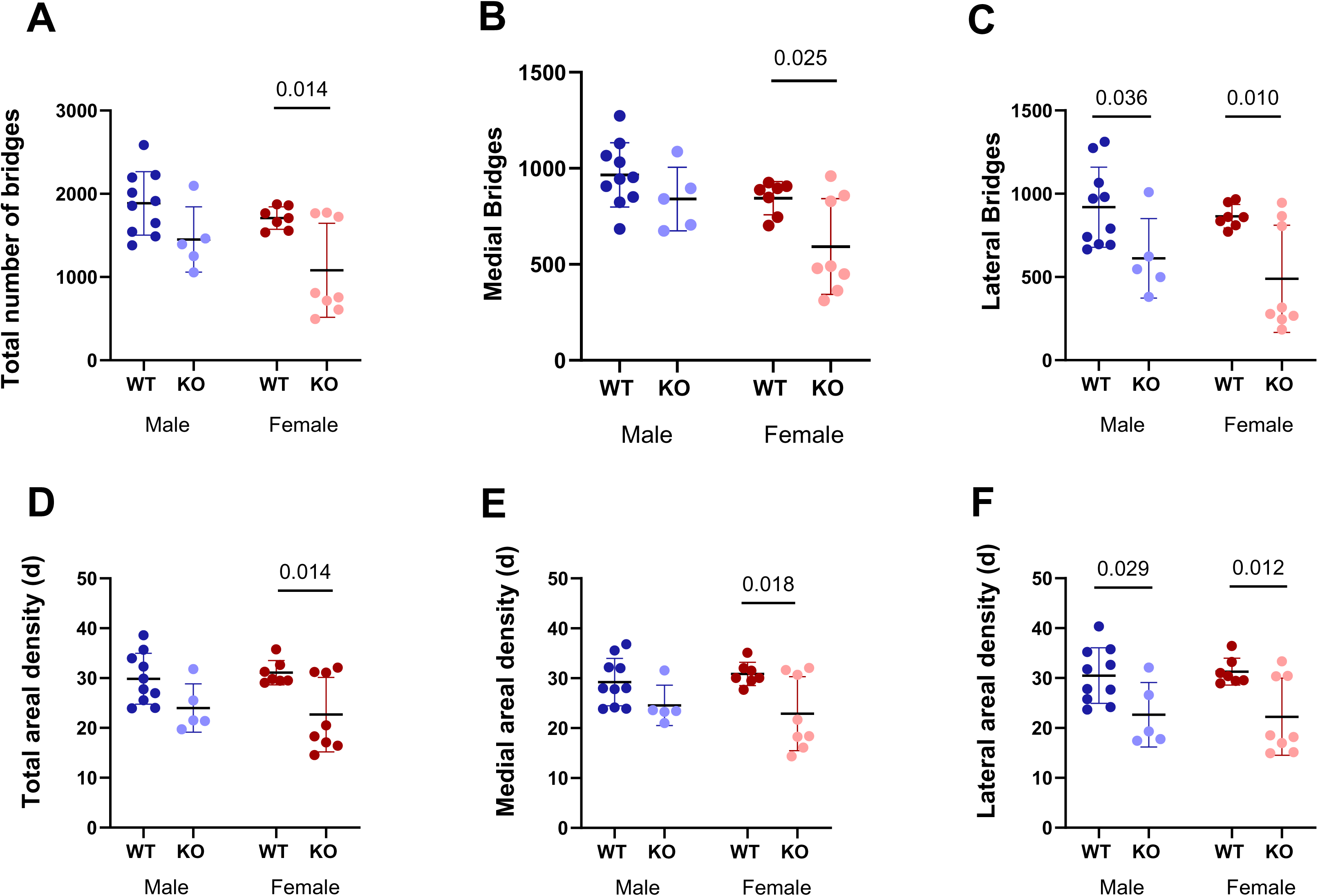
Growth plate bony bridges are reduced in old female but not male TRAP^-/-^ mice. A) Total number of bridges, B) number of medial bridges, C) number of lateral bridges, and D) total, E) medial and F) lateral areal densities of bridges in tibiae of WT and TRAP^-/-^, male and female mice. Areal density is defined as the number of bridges per 256 μm × 256 μm window in each knee joint. Data are presented as individual points with mean ± SD. Two-way ANOVA followed by Tukey’s multiple comparison test with *p*-values for WT vs-KO indicated in the graph. *N* = 5–10 per sex and genotype.

No differences were detected in number or areal density of epiphyseal bony bridges between old WT female and male mice, or between old TRAP^-/-^ female and male mice.

### 3.6 Absence of cortical loading response in old male and female TRAP-/- mice

Our previous study demonstrated absence of cortical loading response in young adult TRAP^-/-^ male mice ^5^. To investigate the bone-loading response in old genotypes, a 2-week repeated loading protocol was applied based on *ex-vivo* pre-gauging strain analyses, as described in the methods section and Supplementary figure 1. In WT mice, loading induced an expected significant increase in total cross-sectional area and cortical area in both males and females (Supplementary table 1), with no loading-responses registered in marrow area or cortical BMD (Fig. 5A-D). In TRAP^-/-^ mice however, no loading-responses were detected in any of the cortical bone parameters, in neither male nor female mice (Supplementary table 1). Comparisons of load-induced fold-changes in cortical parameters demonstrated a significantly greater response in cortical area in both WT males (1.4-fold) and WT females (1.3-fold), compared to their TRAP^-/-^ littermates (Fig. 5B). For female mice, a significantly greater response of 1.2-fold was also detected in the total cross-sectional area (Fig. 5A). There were no intersexual differences in cortical loading responses, independent of genotype.

**Figure 5.**
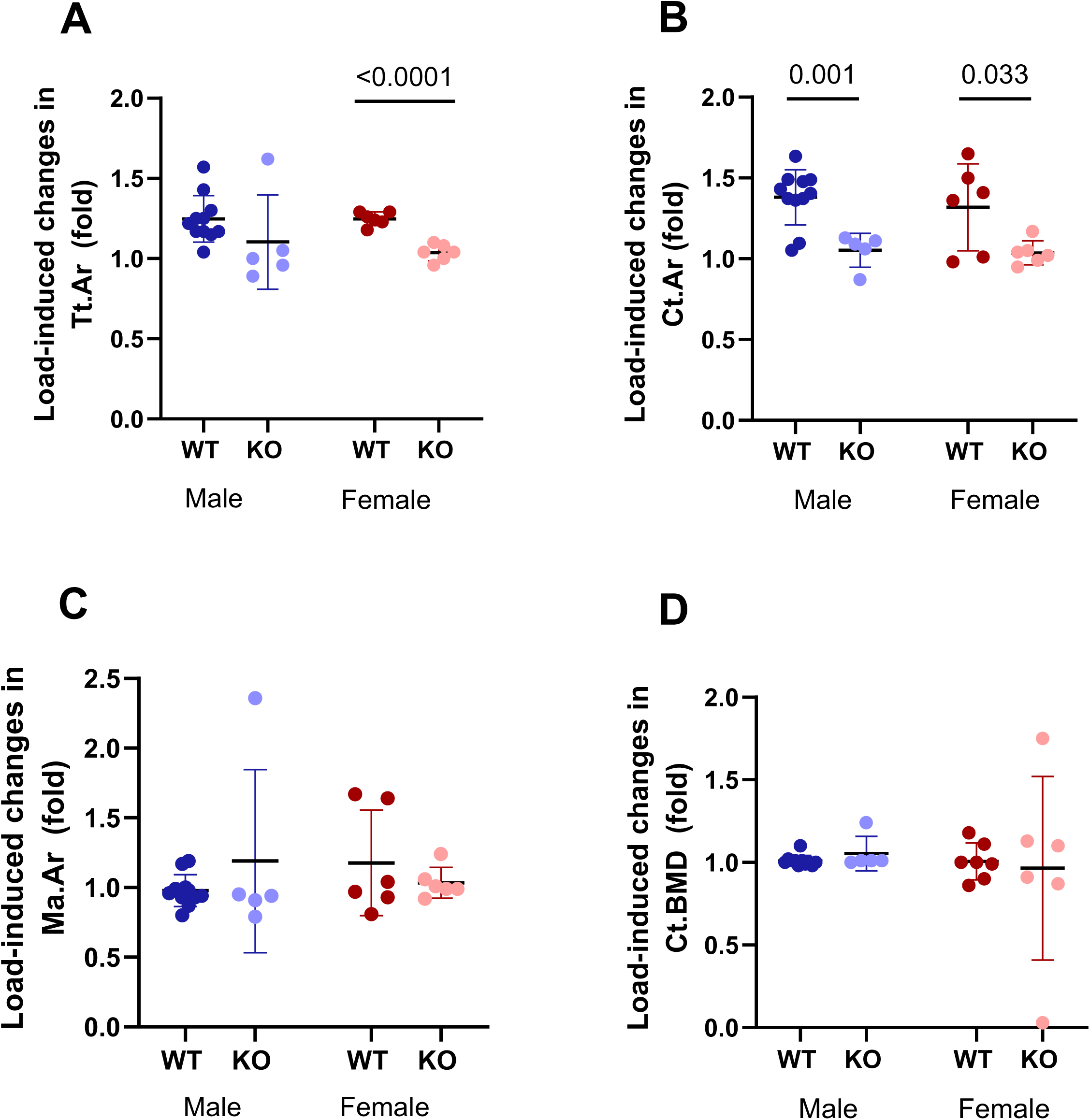
Absence of load-induced cortical bone formation in TRAP^-/-^ mice. A) Periosteally enclosed area (Tt.Ar), B) cortical area (Ct.Ar) and C) marrow area (Ma.Ar), and D) cortical BMD. Load-induced change is expressed as fold-change in loaded leg compared to unloaded leg. Data are presented as individual points with mean ± SD. Two-way ANOVA followed by Tukey’s multiple comparison test with *p*-values for WT vs-KO indicated in the graph. *N* = 5–10 per sex and genotype.

### 3.7 Absence of trabecular loading response in old male and female TRAP^-/-^ mice

We next examined how mechanical loading influences trabecular bone in old male and female WT and TRAP^-/-^ mice (Fig. 6). In old WT mice, both males and females demonstrated the expected increase in trabecular bone parameters in response to loading, with significant increase in trabecular bone volume fraction, trabecular thickness, and trabecular number (Supplementary table 1)(Fig. 6A-C). In contrast, there was a complete absence of any loading response in trabecular bone parameters in both male as well as female TRAP^-/-^ (Supplementary table 1). When comparing the load-induced fold-changes, male WT mice exhibited a significantly greater response of +2.3-fold increase in trabecular bone volume fraction in response to mechanical loading, compared to a +1.3-fold increase TRAP^-/-^ males (Fig. 6A). In females, these differences were even more pronounced, as WT female mice displayed significantly greater response to mechanical loading in trabecular bone volume fraction, trabecular thickness, and trabecular number, with increases of 3.3-fold, 1.4-fold and 4.8-fold, respectively, than female TRAP^-/-^ (Fig. 6A-C). No differences in trabecular loading response between males and females, regardless of genotype.

**Figure 6.**
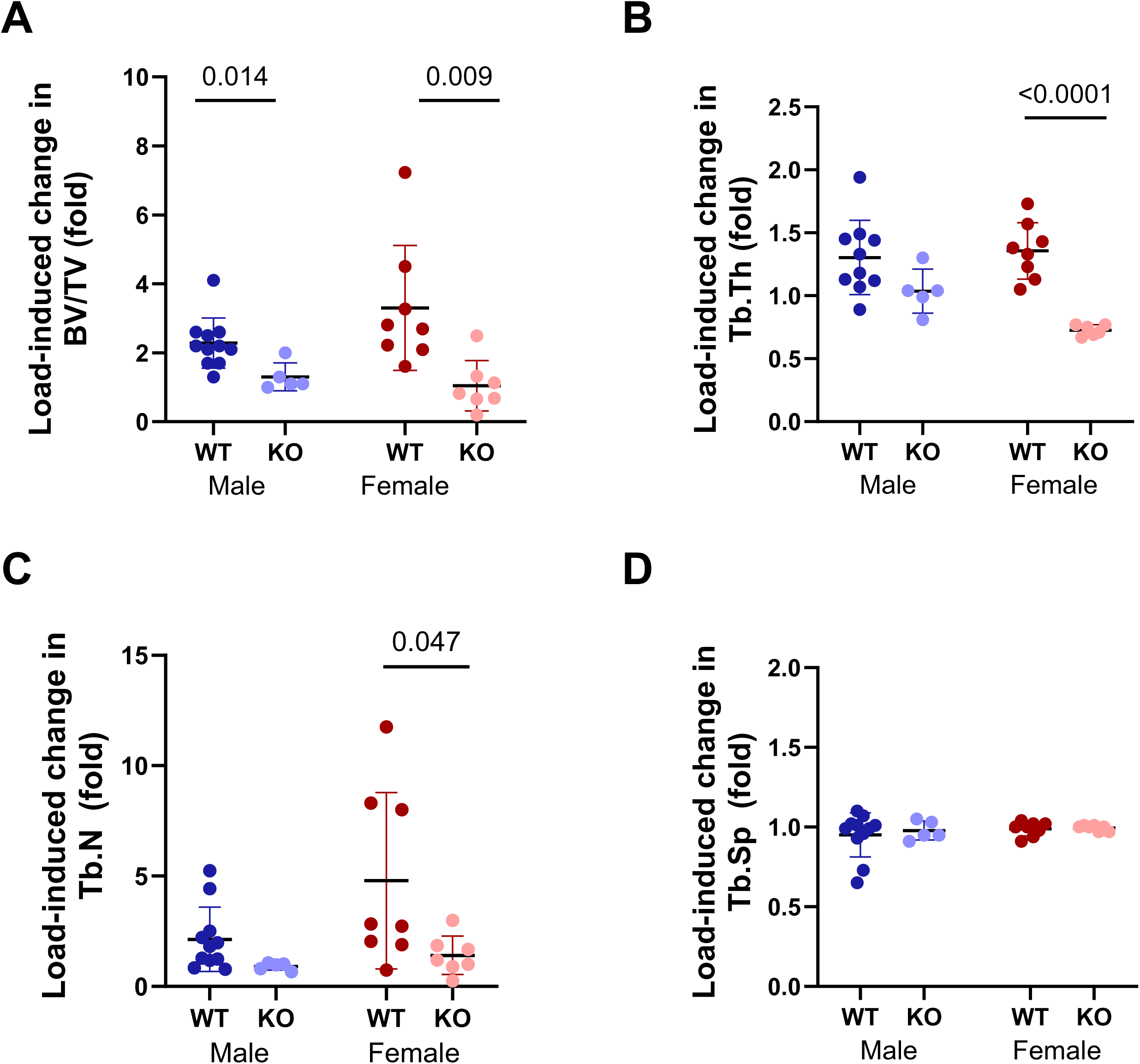
Absence of load-induced trabecular bone formation in TRAP^-/-^ mice. (A) trabecular bone volume fraction (BV/TV), (B) trabecular thickness (Tb.Th), (C) trabecular number (Tb.N), and (E) trabecular separation (Tb.Sp). Load-induced change is expressed as fold-change in loaded leg compared to unloaded leg. Data are presented as individual points with mean ± SD. Two-way ANOVA followed by Tukey’s multiple comparison test with *p*-values for WT vs-KO indicated in the graph. *N* = 5–10 per sex and genotype.

### 3.8 Absence of load-induced response in growth plate bridges in old mice, independent of TRAP

In young WT male mice, we previously found a significant loading-response in number of epiphyseal growth plate bony bridges ^5^. In the current investigation of the effects of TRAP in old male and female mouse bone, we therefore investigated growth plate bony bridges and their areal densities in response to loading. There were no loading-responses registered in neither number nor areal density of bony bridges in old mice, regardless of sex and genotype (Fig. 7A-F).

**Figure 7.**
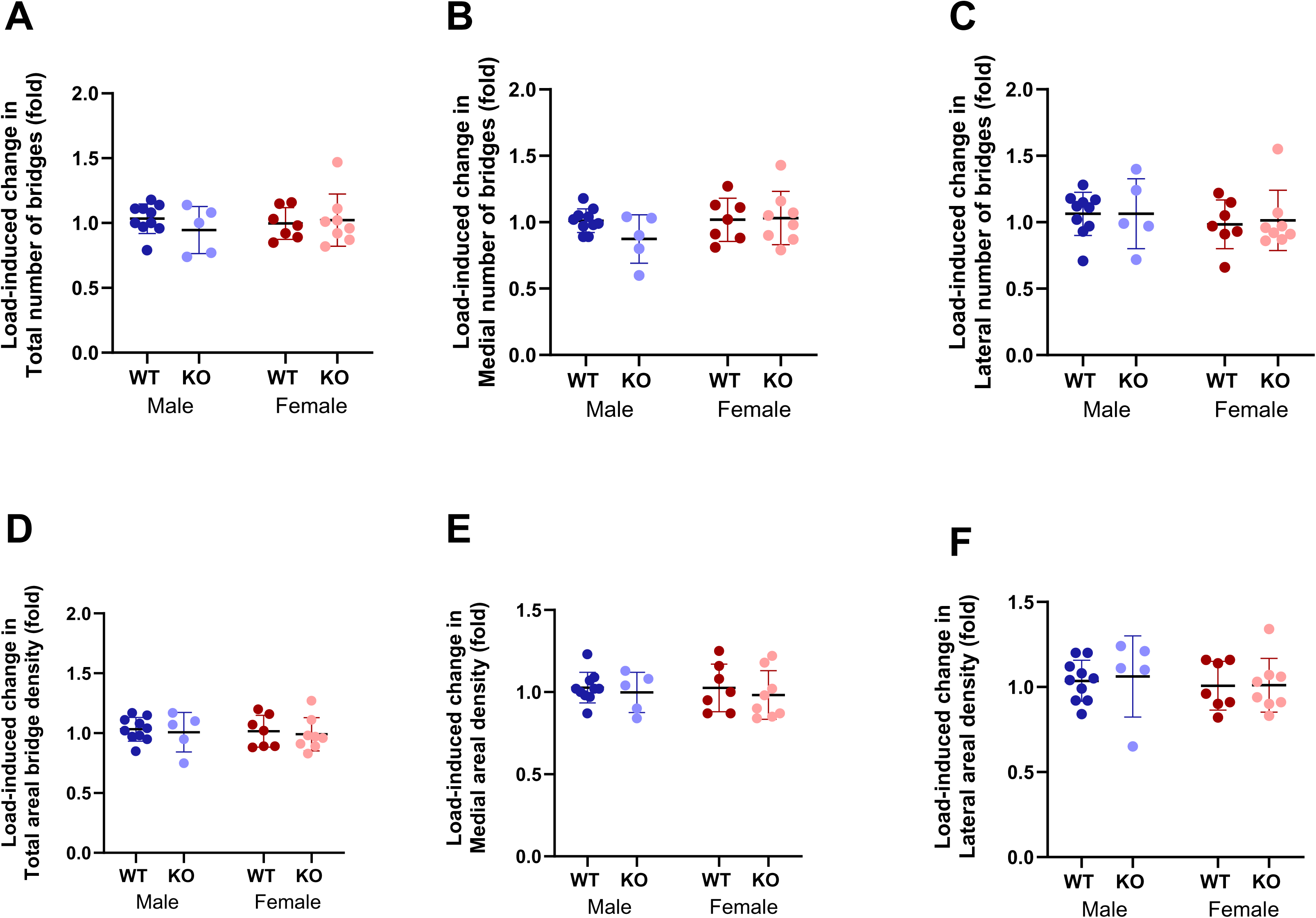
Absence of load-induced response in growth plate bridges in old mice. A) Total number of bridges, B) number of medial bridges, C) number of lateral bridges, and D) total, E) medial and F) lateral areal densities of bridges in tibiae of WT and TRAP^-/-^, male and female mice. Load-induced change is expressed as fold-change in loaded leg compared to unloaded leg. Data are presented as individual points with mean ± SD. Two-way ANOVA followed by Tukey’s multiple comparison test with *p*-values for WT vs-KO indicated in the graph. *N* = 5–10 per sex and genotype.

### 3.9 The *in vitro* loading response in TRAP^-/-^ cells is partially sexually dimorphic

To investigate the responses to mechanotransduction at the cellular level, hematopoietic progenitor cells from the old mice were subjected to *in vitro* loading assays, according to previously established protocols ^18^. In response to high-but not low-intensity *in vitro* loading, osteoclast formation was enhanced from hematopoietic progenitor cells of both male and female WT mice. This effect was absent in cells from TRAP^-/-^ mice of both sexes (Fig. 8A). However, when investigating the responses in extracellular ATP, and cell permeability following *in vitro* loading, clear sex-differentiated patterns were observed. While extracellular ATP increased in high-but not in low-intensity *in vitro* loading in condition medium from WT males, no response was observed in condition medium from TRAP^-/-^ males (Fig. 8B), and cell permeability was not affected by *in vitro* loading intensity in cells from neither WT nor TRAP^-/-^ male mice (Fig. 8C). In contrast, high-, but not low-intensity *in vitro* loading increased extracellular ATP in condition medium of both WT and TRAP^-/-^ female mice (Fig. 8B), and both low- and high-intensity *in vitro* loading decreased cell permeability in female WT and TRAP^-/-^ cells (Fig. 8C). Supporting these findings, gene expression analyses of hematopoietic cells from a subset of WT mice demonstrate the significant increase in ACP5 expression under high-intensity loading, suggesting a transcriptional upregulation of TRAP in response to mechanical stimulation (Fig. 8D).

**Figure 8.**
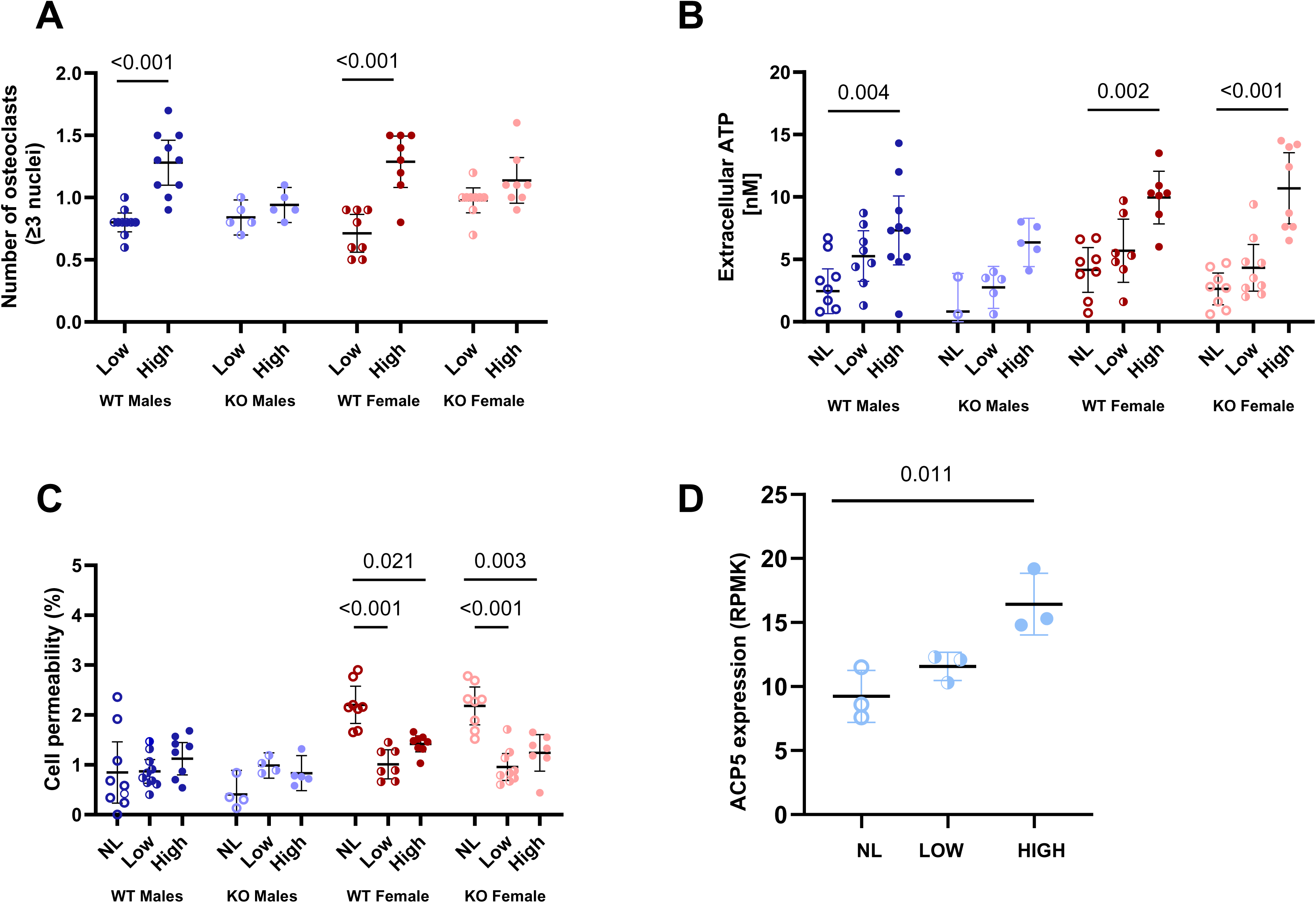
The *in vitro* loading response in TRAP^-/-^ cells is partially sexually dimorphic. (A) Osteoclast formation, (B) extracellular ATP levels, and (C) cell permeability of hematopoietic cells from wild-type (WT) and TRAP^-/-^ (KO) male and female mice. (D) APC5 gene expression in mouse hematopoietic cells in response to *in vitro* loading at low- and high-intensity. Data are presented as individual points with mean ± SD.

## Discussion

This study highlights the pivotal role of TRAP in skeletal architecture and mechanotransduction, with clear sex-related variations in old mice. The study demonstrates significant alterations in bone geometry, cortical and trabecular structure that vary by sex and anatomical site, and deficient mechanical responsiveness in old TRAP^-/-^ mice.

Consistent with our previous research in young adult TRAP^-/-^ mice, old TRAP^-/-^ animals displayed significantly reduced tibial lengths relative their WT littermates, independent of sex. However, we observed no interaction effect for bone length between sexes in old mice, in contrast to the pronounced sex-specific differences in bone length reported earlier in young animals^5^. Our results are in line with previous research showing skeletal alterations characterized by shortened bones, widened and disorganized growth plates, and mild osteopetrosis which have been attributed to impaired osteoclast function and disrupted endochondral ossification during development^4–6^. The levels of the bone resorption marker CTX-1 and the bone formation marker P1NP were significantly lower in old compared to those in young mice of either genotype reported before ^5^. The reduced bone length in old TRAP^-/-^ mice was accompanied with a further reduction in circulating CTX-1 levels but not P1NP. The persistence of reduced bone length into advanced age with the expected age-related decline in systemic turnover markers (CTX-1 and P1NP) and a further reduction in CTX-1 in old TRAP^-/-^ mice suggests that this structural deficit can arise from developmental disturbances and ongoing differences in bone resorption during ageing.

At 19-months of age, TRAP-deficient mice of both sexes remain lighter than their WT littermates, yet only significantly in female mice, and the genotype × sex interaction that characterised the juvenile phenotype is no longer evident. Interestingly, several visceral-organ masses were significantly affected in female TRAP^-/-^ mice, while remaining unaffected in TRAP^-/-^ males. In young adult (12-week-old) TRAP^-/-^ mice, we have reported a larger deficit in body mass and bone length that was markedly greater in males than females, despite normal serum IGF-1 and stable organ masses ^5^. Together, these findings suggest that the primary growth impairment caused by loss of TRAP arises early in life, but that sex-specific amplifiers, potentially androgen-driven differences in bone modelling, diminish with ageing. The persistence of reduced body weight into late adulthood, without parallel changes in liver, spleen, lung or reproductive-organ size in males, argues against a global endocrine defect and instead supports a bone-centred mechanism whereby shorter long bones continue to constrain overall mass in male mice. That most organs scale normally despite lifelong skeletal shortening implies effective physiological compensation, consistent with the mild, non-morbid phenotype observed in human TRAP deficiency^15^. In contrast, the significant changes in internal-organ sizes in old TRAP^-/-^ females indicates more persistent and systemic effects in females. The observed increase in uterine weight also after adjusting to body mass in female TRAP^-/-^ mice compared to WT controls may reflect elevated circulating oestrogen levels in female TRAP^-/-^ mice. Oestrogen is known to promote growth-plate closure, endosteal bone apposition and increase trabecular bone mass^20,21^, suggesting that the skeletal phenotype observed in female TRAP^-/-^ mice could, at least in part, be mediated by altered estrogen levels.

The cortical bone phenotype remains altered in old TRAP^-/-^ males, although less pronounced compared to younger mice. Interestingly, in old mice, also female TRAP^-/-^ displayed altered cortical bone structure parameters that were not detectable in younger mice^5^. More specifically, in TRAP^-/-^ males, the changes were seen as increased cortical cross-sectional area and increased total cortical area, whereas in TRAP^-/-^ females the changes observed were increased cortical BMD and decreased marrow area. Similar cortical thickening observed in younger male TRAP^-/-^ mice has been attributed to impaired osteoclastic resorption at endocortical and periosteal surfaces, leading to increased cortical mass^22^. Our data extend these findings to old mice, reinforcing the notion that cortical bone remodelling may be less susceptible to hormonal variations and more consistently influenced by TRAP activity in male mice. The increased uterine mas, decreased serum CTX-1 and unaffected serum P1NP in old female TRAP^-/-^ mice indicate decreased endosteal bone resorption due to either increased oestrogen levels, or impaired osteoclast function, or both as a mechanism for the decreased marrow area in old female TRAP^-/-^ mice.

Analysis of trabecular bone also revealed marked sex-specific differences in the old mice. Male TRAP^-/-^ mice showed significant increase in trabecular bone compared to WT mice, while there were no effects visible in the female TRAP^-/-^ trabecular bone. The significantly elevated trabecular bone volume fraction, trabecular thickness and trabecular number in TRAP^-/-^ males demonstrate a sustained disruption in trabecular bone previously identified in younger male TRAP^-/-^ mice during ageing^4–6^. In contrast, the lack of changes in old female TRAP^-/-^ mice suggest possible compensatory or protective mechanisms modulated by oestrogen or sex-specific mechanical loading patterns. Supporting our findings in the young adult TRAP*^-/-^* mice, the phenotypic feature in trabecular bone is sustained in old male TRAP^-/-^, while remaining unchanged in old female TRAP^-/-^ mice. Notably though, it seems as the severity of the trabecular phenotype is reduced in old male TRAP^-/-^ mice compared to their younger counterparts. In younger animals, TRAP deficiency is associated with excessive accumulation of trabecular bone due to impaired resorption and delayed remodelling. However, in old TRAP^-/-^ mice, although trabecular volume remains elevated in males, the phenotype is less pronounced. This attenuation could reflect a partial adaptation or compensatory remodelling that occurs over time, possibly through alternative osteoclastic pathways or age-related shifts in bone turnover dynamics. Alternatively, it may suggest a natural age-associated decline in trabecular remodelling in WT animals that narrows the phenotypic gap with TRAP-deficient mice. Sex differences in age-related changes in cortical and trabecular bone morphology, structure, and remodelling is well-documented ^23,24,25^. Our clear sex-related variabilities in old mice supports previous findings ^26^ and further demonstrate the importance of including both sexes also in animal models of bone and skeletal biology.

Analysis of growth plate bridges that spam across the growth plates revealed marked sex-specific differences in the old mice, where the total number and areal density, for both medial and lateral bridges, were reduced in female, but not male, TRAP^-/-^ mice. This contrasts with young TRAP^-/-^ mice where male mice had fewer total number of bridges at both the medial and lateral side, while female TRAP^-/-^ mice had normal number of bridges compared to WT mice^5^. Young TRAP^-/-^ mice of both sexes had unchanged total areal density of bony bridges in the growth plate. Thus, the normal increase in number and areal density of growth plate bridges that spam across the growth plates during ageing is partially disrupted in female, but is normalised in male, TRAP^-/-^ mice. As bridges spanning the growth plate are thought to support the growth plate, one may speculate that, in old female TRAP^-/-^ mice, less bridges are needed due to a narrower marrow space. Studies on bony bridges in old mice are limited, but existing data suggest increased bridging with age, especially in areas of mechanical stress ^27^. We previously found that the number and density of growth plate bridges increase in response to loading in young adult mice ^7^. Interestingly, the number and density of growth plate bridges were not altered by loading in old mice irrespective of genotype or sex. Notably, our study is the first to investigate loading-responses in growth plate bony bridges in old mice.

A critical aspect of our findings is the significant impairment of load-induced anabolic bone responses in TRAP^-/-^ mice. Regardless of the sexual dimorphisms in effects of TRAP on old mice, observed in internal organ sizes, cortical bone parameters, trabecular bone parameters, and growth plate architecture, neither male nor female old TRAP^-/-^ mice showed any anabolic bone response to mechanical loading. The significant responses in the old WT animals confirms that the applied loading protocol has beneficial anabolic effects on bone even in old animals of both sexes. Although standardized mechanical strain was applied, TRAP*^-/-^*bones required higher forces, indicating alterations in bone geometry or matrix composition, potentially increasing stiffness and reducing adaptive responsiveness as previously seen in young mice^5^. This impaired mechano-adaptive capacity aligns with previous studies that show TRAP’s involvement in matrix protein processing, notably osteopontin phosphorylation, essential for osteocyte and osteoblast signalling ^28^. Osteocytes play a central role in bone mechanotransduction through signalling molecules such as sclerostin and extracellular ATP. Loading stimulates ATP release from osteocytes and osteoblasts, activating purinergic signalling pathways critical for osteogenic differentiation and adaptive remodelling^8,29^. TRAP deficiency may indirectly disrupt these osteocyte-driven signalling pathways by altering matrix phosphorylation states, diminishing bone’s sensitivity and responsiveness to mechanical signals. Further investigation is needed to determine precise molecular interactions between TRAP, extracellular matrix modifications, and mechanotransducive pathways.

Comparing phenotypes between young and old TRAP^-/-^ mice highlights critical age-dependent changes, especially in females. While developmental disturbances dominate younger mice as seen in previous publications^4–6^, our models using old mice exhibit functional impairments in mechanically induced bone adaptation. This progression suggests TRAP’s evolving role from facilitating normal growth plate maturation and bone lengthening in youth to maintaining mechanical adaptability in old mice. Disrupted osteocyte signalling due to accumulated, improperly processed extracellular matrix components may become increasingly detrimental with age, manifesting as impaired mechanotransduction and bone rigidity. In addition to its indirect effects on osteocytes and the extracellular matrix, TRAP appears to play a direct role in osteoclast induction. Our *in vitro* data demonstrate that high mechanical loading robustly induces osteoclastogenesis in both male and female WT mice, indicating a consistent mechanoresponsive osteoclastogenic potential across sexes. In contrast, TRAP^-/-^ mice of both sexes failed to exhibit any significant increase in osteoclast formation under similar loading conditions. This impaired osteoclastogenic response aligns with earlier findings that TRAP expression is upregulated during osteoclast differentiation and is required for the proper fusion and activation of osteoclast precursors ^30,31^. TRAP may facilitate the resorptive activity of osteoclasts through modulation of actin ring formation, vesicular trafficking, and production of reactive oxygen species essential for matrix degradation ^32,33^. These sex-independent impairments observed in TRAP-deficient models suggest that TRAP is indispensable for the mechanosensitive induction of osteoclasts. Therefore, in the absence of TRAP, mechanical cues are insufficient to activate the full osteoclastogenic program, leading to compromised bone remodelling in both males and females. These findings reinforce TRAP’s critical role as a mediator of osteoclast function and highlight its necessity for translating mechanical stimuli into cellular remodelling responses.

In conclusion, TRAP deficiency substantially compromises skeletal integrity, and mechanical adaptability across the lifespan, with age- and sex-specific consequences. The clear sex-related variability in skeletal phenotype and bone maintenance seen in old TRAP^-/-^ mice, combined with the complete absence of anabolic responses to loading in absence of TRAP, in both males and females, underscores TRAP’s complex roles beyond osteoclast function alone, highlighting critical interactions between osteoclast-mediated remodelling, osteocyte signalling, and matrix biology. Future studies should explore therapeutic strategies targeting these interactions, particularly addressing matrix phosphorylation and osteocyte signalling pathways, to preserve skeletal integrity and adaptive capacity in aging and metabolic bone diseases.

## Acknowledgements

This study was supported by the Swedish Research Council grant number 2019-01295 (SW), and Swedish Cancer Society grant number 190165 (GA). We thank Dr. Gabriel G Galea, UCL, UK, for technical assistance with the SSA analysis. We thank Dr. Sakshi Vats, Dr. Frank Ning, Dr. Dorina Ujvari, Joshua Sweidan, and Johan Lännerström for assistance with experiments.

