## Supplementary Table for "Tartrate-resistant acid phosphatase (TRAP/ACP5) sex-specifically regulates bone maintenance in old mice, but the anabolic effects of mechanical loading is regulated in a sex-independent way"

### Supplementary Table 1

| Supplementary table 1. Loading-responses in bone parameters in WT and TRAP -/- males and females | | | | | | | | | | | | |
|---|---|---|---|---|---|---|---|---|---|---|---|---|
| | Males | | | | | | Females | | | | | |
| | WT | | | TRAP -/- | | | WT | | | TRAP -/- | | |
| **Bone parameters** | **Non-loaded** | **Loaded** | ***P*** | **Non-loaded** | **Loaded** | ***p*** | **Non-loaded** | **Loaded** | ***p*** | **Non-loaded** | **Loaded** | ***p*** |
| Tt.Ar | 1.87 ± 0.25 | 2.31 ± 0.17 | **<0.0001** | 2.43 ± 0.53 | 2.59 ± 0.26 | 0.052 | 1.27 ± 0.11 | 1.58 ±0.09 | **<0.0001** | 1.22 ± 0.12 | 1.27 ± 0.13 | 0.105 |
| Ct.Ar | 1.06 ± 0.10 | 1.45 ± 0.12 | **<0.0001** | 1.34 ± 0.28 | 1.40 ± 0.29 | 0.654 | 0.73 ± 0.09 | 0.95 ± 0.16 | **0.005** | 0.85 ± 0.14 | 0.89 ± 0.15 | 0.830 |
| Ma.Ar | 0.88 ± 0.15 | 0.85 ± 0.14 | 0.387 | 1.10 ± 0.29 | 1.19 ± 0.32 | 0.699 | 0.54 ± 0.07 | 0.63 ± 0.19 | 0.311 | 0.37 ± 0.05 | 0.38 ± 0.07 | 0.435 |
| Ct.BMD | 2.44 ± 0.02 | 2.46 ± 0.08 | 0.323 | 2.35 ± 0.20 | 2.45 ± 0.01 | 0.320 | 1.38 ± 0.09 | 1.40 ± 0.17 | 0.771 | 1.60 ± 0.25 | 1.45 ± 0.72 | 0.692 |
| BV/TV | 4.56 ± 1.32 | 9.76 ± 1.88 | **<0.0001** | 10.06 ± 7.59 | 11.46 ± 7.30 | 0.182 | 2.96 ± 1.63 | 7.54 ±2.15 | **<0.0001** | 3.73 ± 2.77 | 3.13 ± 2.37 | 0.720 |
| Tb.Th | 0.055 ± 0.010 | 0.069 ± 0.010 | **0.004** | 0.070 ± 0.010 | 0.072 ± 0.010 | 0.067 | 0.062 ±0.010 | 0.082 ± 0.010 | **0.002** | 0.071 ± 0.030 | 0.057 ± 0.020 | 0.038 |
| Tb.N | 0.78 ± 0.26 | 1.39 ± 0.49 | **0.004** | 1.58 ± 0.75 | 1.44 ± 0.81 | 0.812 | 0.47 ± 0.24 | 1.47 ± 0.46 | **0.0002** | 0.42 ± 0.19 | 0.43 ± 0.21 | 0.999 |
| Tb.Sp | 0.25 ± 0.01 | 0.26 ± 0.01 | 0.667 | 0.25 ± 0.02 | 0.25 ± 0.02 | 0.403 | 0.27 ± 0.01 | 0.27± 0.01 | 0.462 | 0.27 ± 0.01 | 0.27± 0.01 | 0.547 |
| Tot N bridges | 1885 ± 381 | 1950 ± 486 | 0.432 | 1452 ± 392 | 1387 ± 563 | 0.632 | 1709 ± 135 | 1697 ± 155 | 0.875 | 1081 ± 766 | 1060 ± 509 | 0.759 |
| Medial N bridges | 966 ± 168 | 980 ± 222 | 0.661 | 840 ± 166 | 736 ± 233 | 0.614 | 844 ± 87 | 852± 106 | 0.875 | 592 ± 249 | 587 ± 210 | 0.898 |
| Lateral N bridges | 919 ± 241 | 970 ± 271 | 0.384 | 612 ± 239 | 651 ± 336 | 0.055 | 865 ± 71 | 844 ± 134 | 0.753 | 489 ± 322 | 473 ± 301 | 0.572 |
| Tot areal density | 30.0 ± 5.1 | 30.9 ± 6.7 | 0.356 | 24.0 ± 4.9 | 24.1 ± 6.3 | 0.951 | 31.1 ± 2.4 | 31.5 ± 3.9 | 0.782 | 22.7 ± 7.5 | 22.1 ± 7.1 | 0.621 |
| Medial areal density | 29.2 ± 4.7 | 30.1 ± 6.4 | 0.405 | 24.6 ± 4.1 | 24.6 ± 5.7 | 0.991 | 30.9 ± 2.4 | 31.6 ± 4.2 | 0.673 | 22.9 ± 7.4 | 22.2 ± 7.3 | 0.619 |
| Lateral areal density | 30.5 ± 5.6 | 31.6 ± 7.2 | 0.393 | 22.7 ± 6.5 | 23.5 ± 7.1 | 0.763 | 31.3 ± 2.7 | 31.3 ± 3.9 | 0.986 | 22.2 ± 7.7 | 22.0 ± 6.9 | 0.824 |
| Statistical analyses conducted through repeated measures two-way ANOVA, followed by paired T-test for pairwise comparisons | | | | | | | | | | | | |

### Supplementary Figure 1.

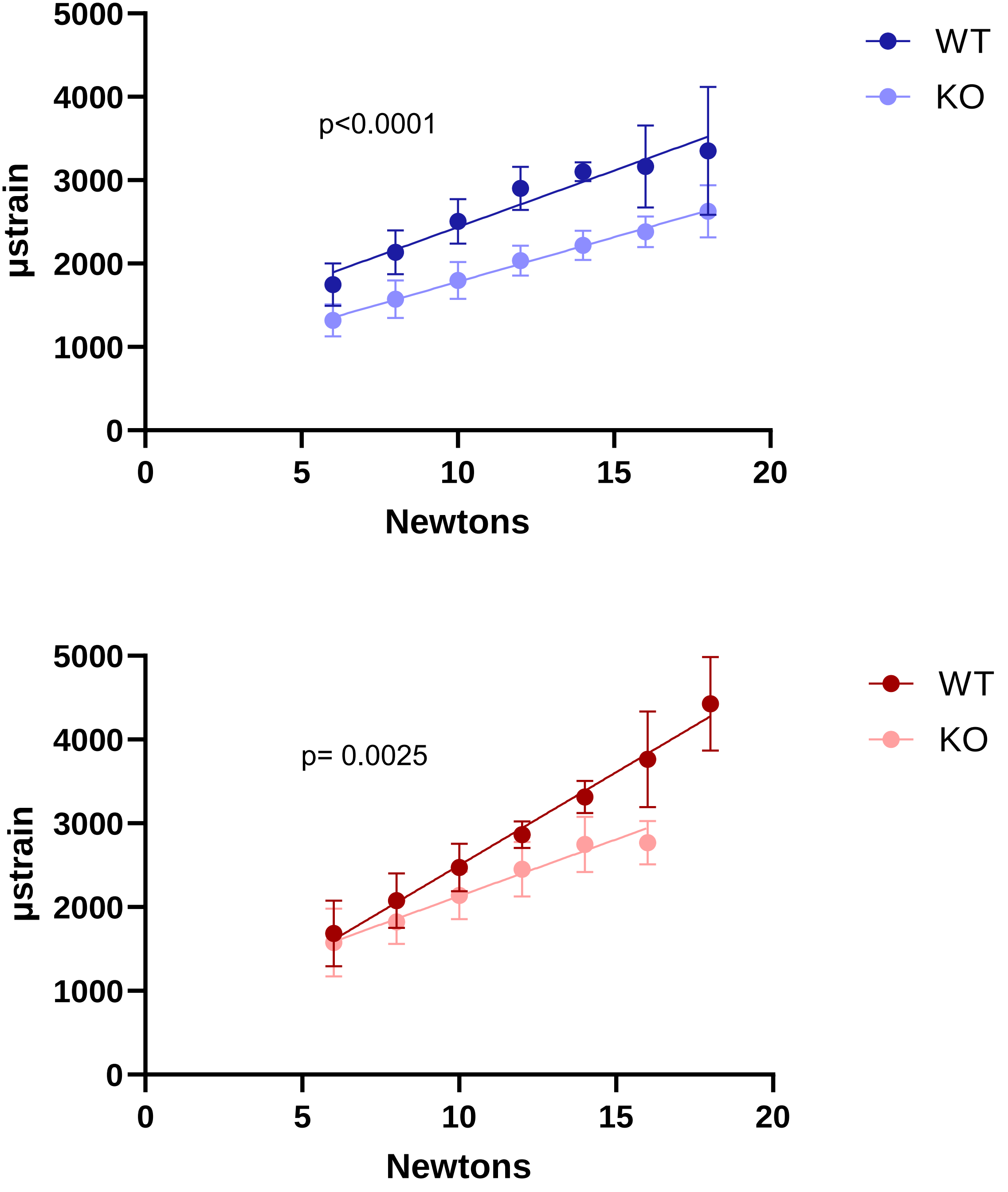
